# First report on *Tropilaelaps mercedesae* presence in Türkiye and insight into potential westward invasion corridors

**DOI:** 10.64898/2026.09.15.751928

**Authors:** Aslı Özkırım, Aleksandar Uzunov, Cecilia Costa, İrfan Kandemir

**Affiliations:** Faculty of Science, Department of Biology, Hacettepe University, Beytepe/Ankara, Türkiye; Ss. Cyril and Methodius University in Skopje, Faculty of Agricultural Sciences and Food, Skopje, N. Macedonia; Macedonian Academy of Sciences and Arts, Research Center for Environment and Materials, Skopje, N. Macedonia; State Key Laboratory of Resource Insects, Institute of Apicultural Research, Chinese Academy of Agricultural Sciences, Beijing, China; Consiglio per la ricerca in agricoltura e l’analisi dell’economia agraria (CREA), Centro di ricerca Agricoltura e Ambiente, Bologna, Italy; Department of Biology, Ankara University, Ankara, Türkiye

## Abstract

The honey bee ectoparasitic mite *Tropilaelaps mercedesae* is an emerging threat to Western beekeeping, as it is expanding its range from native Asia. In this study, the first record of the presence of *Tropilaelaps mercedesae* in Türkiye, in the Artvin province, in the north-east of the country is documented. Following reports of weakening colonies, eight apiaries were inspected using the rapid brood decapping method. All apiaries were found positive, and infestation was detected in 22 of 24 examined colonies. Mite samples were analysed with morphometry and barcoding and were confirmed as *T. mercedesae*. In the paper, we discuss the impact of imports and migration practices on the mite’s occurrence in Türkiye. In addition, we forecast the potential corridors and means of further invasion and address the implications for European and global apiculture.

## Introduction

Originally hosted by the open-nesting, single-comb Asian giant honey bees (*A. dorsata* and *A. laboriosa*), the ectoparasitic mite *Tropilaelaps mercedesae* shifted to the Western honey bee (*Apis mellifera*) after the latter’s introduction to Asia around a century ago, becoming one of the major threats to Asian beekeeping (Chantawannakul et al., 2018).

After having stayed confined to Asia since its first description as a Western honey bee parasite (Delfinado, 1963), a westward expansion of the mite into Central Asia and Eastern Europe is currently ongoing (Brandorf et al., 2025; Janashia et al., 2024) and has drawn global attention from the beekeeping and scientific communities. This is justified by the mite’s exceptional population growth potential (Aurell et al., 2026) and pathogenicity (Chantawannakul et al., 2018), with devastating consequences on the health of *A. mellifera* colonies. However, almost two years after the initial confirmation from southwest Russia (Krasnodar and Rostov regions) and Georgia, no further incidences have been reported on the European continent, except for an unconfirmed report from Belarus (Franco et al., 2025).

Given the terrestrial border of about 270 km between Georgia and Türkiye, an imminent invasion of the Turkish neighbouring region was anticipated (Janashia et al., 2024). This scenario was considered plausible due to: a) the natural spread of the mite via foraging (Tokach et al., 2025) and swarming (Uzunov et al., 2026) and b) possible cross-border exchange of reproductive material, such as queens which is known to occur, despite being illegal. Thus, systematic monitoring of apiaries for the presence of the mite in the north-eastern part of Türkiye, close to the Georgian border, has been conducted since 2024. This targeted surveillance has now resulted in the first detection of *T. mercedesae* in Türkiye. Here, we report this first detection in the north-eastern province of Artvin and assess the possible extent of the invasion. We further discuss its implications for European and global beekeeping by identifying potential corridors and mechanisms for further spread. Given Türkiye’s strategic geographic and trade position at the intersection of Asia, Europe, and Africa, and the magnitude of its beekeeping capacity, this report has very high implications for global beekeeping biosecurity.

## Methods

Following the invasion of *T. mercedesae* into neighbouring Georgia (Janashia et al., 2024), systematic field monitoring for mite detection was conducted in the Artvin province each spring and autumn from 2024 to 2026 (spring). Location and apiary selection criteria were: a) proximity to the Georgian border line, b) apiary size of more than 40 colonies, and c) beekeeper willingness to collaborate. Apiaries belonged to both stationary and migratory beekeeping practices. Beekeepers were informed via dedicated seminars and were interviewed for information on most recent seasonal migratory operations. Each year, 40-50 apiaries were inspected, and in each apiary 20 - 30 % of the colonies were screened for *T. mercedesae* presence using Rapid Brood Decapping (RBD) described by Uzunov et al. (2025). Briefly, the method relies on the use of wax depilation strips to uncap sealed brood and the fact that *T. mercedesae* promptly exits the cells after disturbance and moves fast across the comb. Brood combs are rapidly inspected directly in the field, and the procedure can also be video-recorded for further visual inspection.

On 26th July 2026, upon request from the Artvin Beekeepers’ Association, which had noticed unusual colony declines and formic acid use by beekeepers, a follow-up screening using the above-noted methodology was conducted in 8 stationary apiaries in the Artvin province (Fig. 1, Tab. 1). In each apiary, on the basis of population decrease, 3 colonies were inspected, resulting in examination of a total of 52 frames from 24 colonies. When colonies were found positive with RBD, individual pupae were extracted from 7 - 10 brood cells using forceps to check for the presence of *T. mercedesae* and *Varroa destructor*. Individual mites were collected and placed in Eppendorf tubes (N=10) containing 99% ethanol and transported to the laboratory. In addition, to collect a higher number of mites, clean wax strips were applied to the decapped comb surface with running mites. Then, the wax strips (N=10) were folded with the adhesive surface facing inward to prevent the samples from drying and were preserved in a sugar-based gel medium. The samples preserved in Eppendorf tubes and the wax strips were marked to identify the date, location, and colony number and transported to the laboratories for identification. to be subjected to morphometric analysis. The individual mites in six (N=6) of the tubes were morphometrically analysed using Olympus BX51 microscope, Olympus SC50 Camera system, cellSens Imaging Software V3 at the Laboratory of MicroBeeotic of Hacettepe University in Ankara, following the method included in the standard diagnostic guidelines of the World Organisation for Animal Health (2018). The mite samples collected on sticky strips were delivered, together with N=4 tubes each with 2-5 individual mites to the Insect Molecular Systematics Laboratory in the Department of Biology at Ankara University. A total of 10 pools of 2-10 mites and mite parts collected from the sticky sheets was used to extract total nucleic acids with the CTAB protocol following Doyle and Doyle (1987). For species delimitation, the cytochrome oxidase subunit I (COI) region of mtDNA was partially amplified using universal primers LCO1490 and HCO2198 (Folmer et al., 1994). The amplification procedure was performed according to Selis et al. (2024). The amplified fragments were visualized on a 1.5% agarose gel, and the both way sequencing was done by the BM Labosis company using forward and reverse primers.

**Figure 1.**
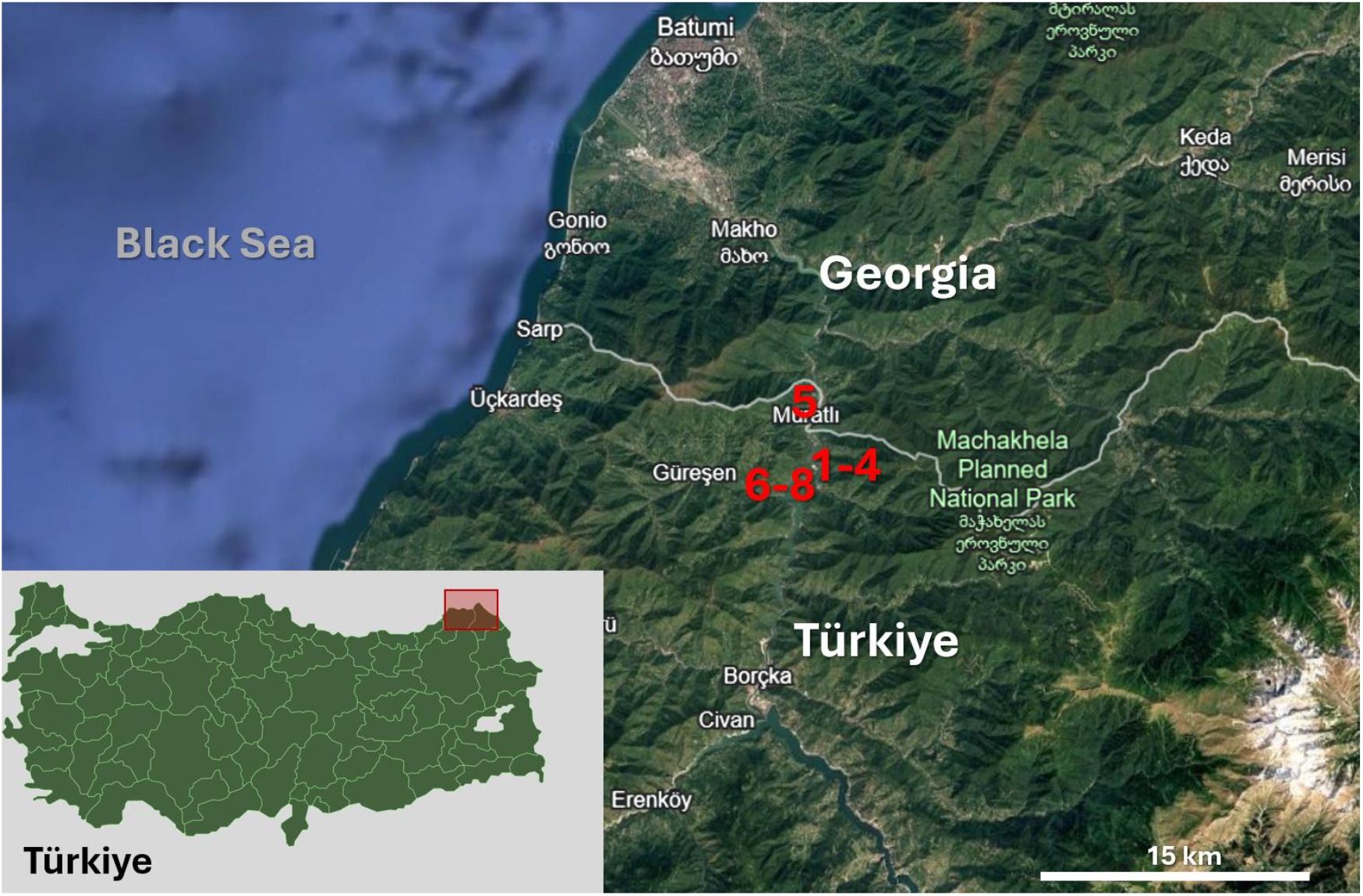
Apiaries’ locations (Google Earth®). Red numbers indicate *T. mercedesae*-inspected and infested apiaries. For more details, see Table 1 with corresponding apiary numbers.

## Results

The inspections performed in 2024 (autumn), 2025 (spring and autumn), and spring 2026 did not reveal the presence of *T. mercedesae*. Instead, the inspections performed in July 2026 (Fig. 1) showed that 22 (91.6%) of the 24 inspected colonies were *T. mercedesae*-positive (Tab. 1). Individual inspection of capped brood cells always confirmed the results obtained by RBD. All infested colonies exhibited weakening, a decrease in the adult population, and deformed adult bees unable to emerge from brood cells.

**Table 1.** Number of inspected apiaries, date, region, location, number of colonies in the apiary (April 2026, July 2026), number of inspected and positive colonies for *T. mercedesae* infestation.

| Api<br>ary<br>N° | Date of<br>inspection<br>(2026) | Region | Location | No. of total<br>colonies<br>April 2026 | No. of total<br>colonies<br>July 2026 | No. of<br>inspected/<br>infested<br>colonies |
| --- | --- | --- | --- | --- | --- | --- |
| 1 | 26.07.26 | Artvin-Borçka | Muratlı-Karşıköy | 100 | 63 | 3/3 |
| 2 |  |  |  | 80 | 42 | 3/2 |
| 3 |  |  |  | 140 | 55 | 3/3 |
| 4 |  |  |  | 50 | 22 | 3/3 |
| 5 |  |  | Muratlı | 70 | 14 | 3/2 |
| 6 |  |  | Çavuşlu | 39 | 25 | 3/3 |
| 7 |  |  | Çavuşlu | 40 | 30 | 3/3 |
| 8 |  |  | Çavuşlu-Sirt Mahalle | 60 | 40 | 3/3 |

The infested colonies were located in a range of 200 m - 40 km south of the border with Georgia (Fig. 1). In all *T. mercedesae* - positive colonies, co-infestation with *V. destructor* was confirmed by inspection of individual brood cells.

Seven of the 10 pools successfully amplified the barcoding region and were sent for sequencing. The obtained sequences (all 688 bp and identical) confirmed that all (N=7) samples belong to *T. mercedesae*, and BLAST analysis showed 99.83% basepair similarity with the specimen from Russia (Accession Numbers OR965215.1), 99.73% with Uzbekistan samples (PX136173.1, PX136174.1, PX136176.1, PX136177.1, PX136552.1, PX136553.1, PX136554.1), 99.72% with Pakistan, Vietnam and Indonesia (LC474398.1, LC474401.1, and LC474405.1 respectively), 99.71% with samples from India (OK188792.1, MW337212.1, and MW337213.1). In addition, the morphometric measurements of the samples (N=6, Tab. 2, Supp. S1), width: 0.5 – 0.6 mm and length: 0.95 – 1.05 mm, are consistent with those previously reported for *T. mercedesae* (Anderson and Roberts, 2013; WOAH Terrestrial Manual, 2018; Janashia et al., 2024).

Finally, the official notification procedure was completed according to national legislation and resulted in official inspections and confirmation of positive Tropilaelaps cases to WOAH (31st August 2026, WOAH/WAHIS report: https://wahis.woah.org/#/in-event/7792/dashboard).

**Table 2.** Morphometric results of the sampled *T. mercedesae* mites.

| No. of sample | No. of Apiary (See Table 1.) | Body Width in mm | Body Length in mm | Genital Length in mm (Epigynial Shield) | Anal Length in mm (Ventrional Shield) |
| --- | --- | --- | --- | --- | --- |
| 1 | 3 | 0.496 | 0.96 | 0.29 | 0.14 |
| 2 | 3 | 0.520 | 0.98 | 0.29 | 0.15 |
| 3 | 4 | 0.516 | 0.97 | 0.31 | 0.13 |
| 4 | 4 | 0.513 | 0.97 | 0.28 | 0.16 |
| 5 | 5 | 0.539 | 0.99 | 0.30 | 0.15 |
| 6 | 8 | 0.497 | 0.99 | 0.31 | 0.14 |

## Discussion

The expansion of *T. mercedesae* range to Türkiye risks having an enormous impact on regional and global apiculture, particularly given the country’s position at the intersection of Asia, Europe, and Africa, which makes it strategically important for apicultural biosecurity. Here, we present the different types of risk posed by the present finding of *T. mercedesae* in the North-East of Türkiye and the potential westward and southward expansion routes.

### A.m. caucasica *conservation area*

The Artvin and Ardahan provinces have been designated since 2000 as an *in situ* conservation area for the protection of *Apis mellifera caucasica*, one of the autochthonous honey bee subspecies of Türkiye. The conservation model rests on geographic isolation and management of colony movements, with the aim of preventing hybridization with non-local bees (principally *A. m. ligustica, A. m. carnica*, and Buckfast origin) that are present elsewhere in Türkiye (Presidential Decree No. 1 on the Organization, Duties and Powers of the Ministry of Agriculture and Forestry, published in the Official Gazette No. 30474 from 10th July 2018). Thus, the introduction of any live honey bees (including queens, nucs, swarms, etc.) from outside the conservation area as well as the movement of colonies from the conservation area to other regions, is prohibited. In contrast, migratory beekeeping is permitted between other provinces of Türkiye that are not designated as *A. m. caucasica* conservation areas and is widely practised throughout the country (Fig. 2). The isolation-based governance was not designed for sanitary reasons; however, given no movement of colonies in and out of the area, it may offer some barrier to further expansion of *T. mercedesae* within Türkiye. Nevertheless, the presence of the mite in this specific area poses an additional threat to the population of *A. m. caucasica*.

**Figure 2.**
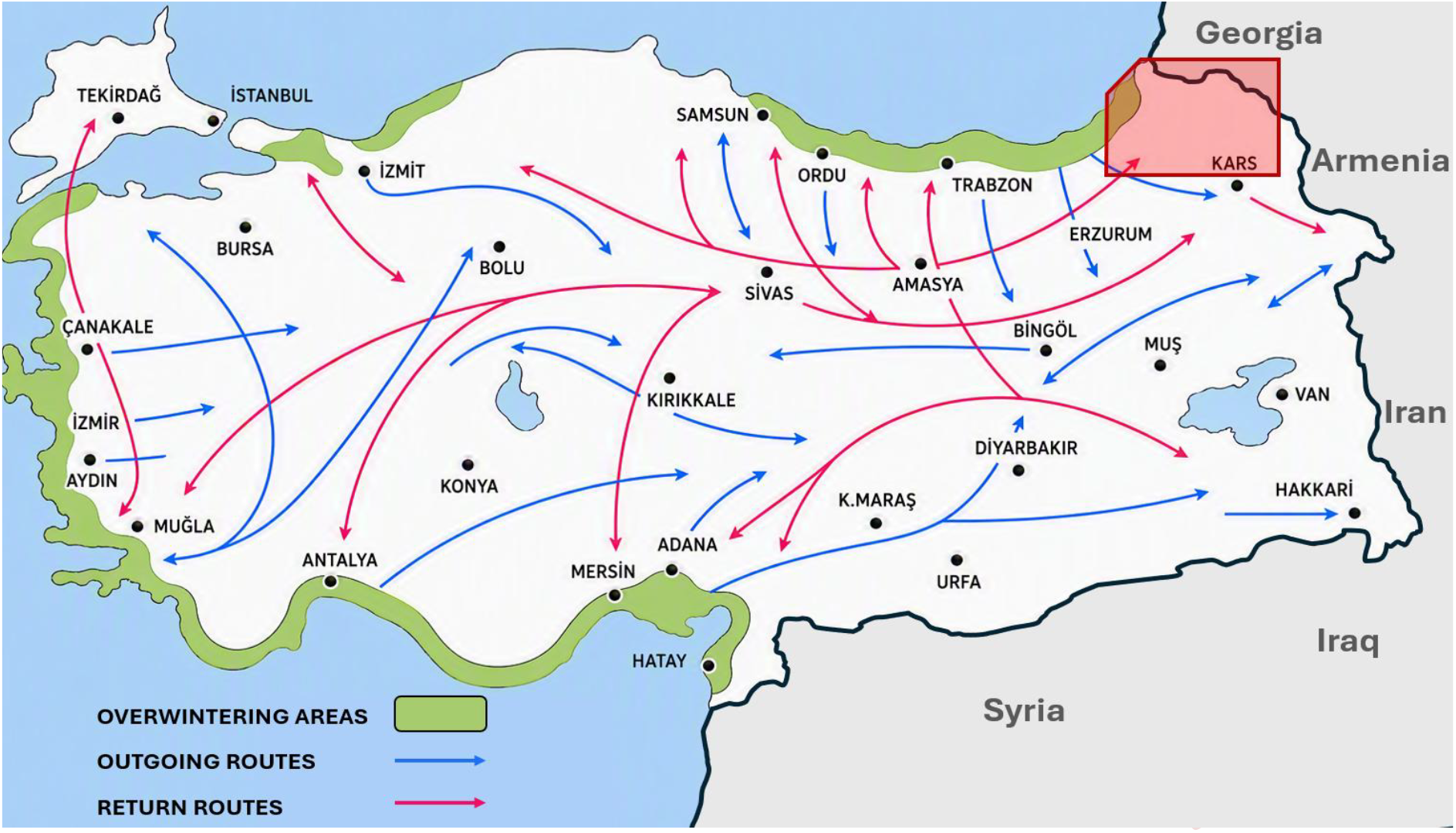
A historical map of common honey bee colony migration routes and overwintering areas in Türkiye (improved after Bahri Yılmaz’s version). The red rectangle indicates isolated regions from all other routes for colony movements and the approximate *T. mercedesae* invaded region.

### Impacts of local migration practices and illegal queen imports

A critical epidemiological detail is that some apiaries in northernmost Artvin were moved to a different part of the province and then to the province of Ardahan (also *A. m. caucasica* conservation area) just after the chestnut (*Castanea sativa*) honey was harvested - roughly a month before the detection of the *T. mercedesae* mites occurred. The eight stationary positive apiaries represent the infestation status of the original Artvin–Borçka area, whereas possible migratory positive apiaries may provide a plausible epidemiological link between this focus and other regions of Artvin and parts of Ardahan. This distinction might be particularly important when interpreting the spatial distribution of positive apiaries, because movement-mediated dissemination may result in secondary foci beyond the area in which the infestation was initially detected (Uzunov et al., 2026).

Therefore, migratory apiaries may have contributed to the geographical expansion of *T. mercedesae* from the original Artvin–Borçka focus into other areas of Artvin. The practice of temporarily concentrating colonies from multiple beekeepers around a single flow is functionally equivalent to a temporary “mega-apiary,” and is the same mechanism implicated in the rapid regional spread of the parasitic mite *Varroa destructor* historically; it should be flagged as a priority target for active surveillance and, ideally, pre- and post-movement *T. mercedesae* inspection protocols for specific to migratory circuits in the whole of the country.

Examination of the beekeeper profile in the affected area indicates that numerous reports and complaint petitions have been submitted by the Artvin Beekeepers’ Association (pers. communication) to the Provincial Directorate of Agriculture and Forestry concerning the alleged illegal acquisition of *A. m. caucasica* queen bees from Georgia by local beekeepers, due to the lower cost. Despite *T. mercedesae* being a brood parasite, and traditionally thought to be able to survive only a few days on adult bees, there is recent evidence highlighting that survival without brood may actually be more likely than previously reported (Gill et al., 2025; Ramirez et al., 2025).

Importantly, this practice is not limited to beekeepers from Artvin and Ardahan; beekeepers from the Black Sea and Eastern Anatolia regions are also reported to purchase Georgian queens imported through unofficial channels for economic reasons. In such a scenario, the principal drivers of spread may prove not to be migratory beekeeping of high number of colonies in Türkiye, but rather the unauthorized importation of queen bees from abroad.

### Colonies, routes, diversity

Extrapolating from provincial and regional dispersal to the national scale, Türkiye’s beekeeping capacity is, in itself, a risk factor (Girişgin et al., 2026). As a beekeeping country that holds one of the largest colony populations in the world, Türkiye had 9.2 million beehives according to 2023 Turkish Statistical Institute data, with more recent industry figures citing nearly 98,000 beekeeping enterprises (TUİK, 2025). Migratory beekeeping is not a marginal practice but the operational backbone of the sector: figures in the literature range from about 50% of colonies moved seasonally, contributing roughly 80% of national honey production (Özkırım, 2018). Colonies are routinely relocated over hundreds of kilometres between the Black Sea and northeastern uplands, the Aegean and Mediterranean coasts, Central Anatolia, and Thrace, following successive nectar flows across roughly nine months of the year (Fig. 2). This creates a dense, nationwide network involving colonies belonging to Türkiye’s several recognized subspecies and ecotypes (*A. m. caucasica* in the northeast, *A. m. anatoliaca* across central and eastern Anatolia, *A. m. syriaca* in the southeast, plus numerous local ecotypes in the Thrace, Yığılca, Aegean regions (Kandemir et al., 2000)). However, due to the conservation status, the migratory operations don’t pass through Artvin and Ardahan on the way to Black Sea and Eastern Anatolian flows, so the conservation zones are epidemiologically isolated from the rest of the national migratory network. This may offer the opportunity of limiting the spread of the mite to the rest of Türkiye, if carefully managed.

### Türkiye as a biosecurity crossroads

Türkiye’s transcontinental position converts a national spread risk into an intercontinental one. Regarding the international level of the issue, if *T. mercedesae* is confined to the restricted Artvin region, its spread is expected to be slower. If the mite is present elsewhere in the Black Sea or Eastern Anatolia regions, however, its potential rate of spread may increase considerably.

The implications run in several directions: westward, Türkiye’s land border with Greece and Bulgaria places the mite within a short logistical step of continental European territory; both the overland route through Thrace and the intensive maritime and commercial traffic through Turkish Black Sea, Aegean, and Mediterranean ports represent plausible pathways, whether through beekeeper-mediated movement, legal or informal trade in queen transition, or potentially by beekeeping materials.

Eastward and southward, Türkiye’s long borders with Armenia, Iran, Iraq, and Syria deserve equal attention: they represent additional, currently unmonitored corridors through which the mite could spread onward into the Middle East, in light of the potential scarcity of Tropilaelaps surveillance infrastructure and limited veterinary capacity to respond in several areas of the Middle East.

Nevertheless, within the framework of the present study, considering the current distribution map of the mite (Franco et al., 2025), it may be anticipated that the mite could move westward toward Thrace through the Black Sea corridor then to Bulgaria and Greece, similarly as distribution probably occurred from the native range, via Kazakhstan and Uzbekistan to Russia and then to Georgia (Esmaeily et al., 2026; Janashia et al., 2024; Joharchi & Stolbova, 2024).

Thus, the decisions and actions made now on Türkiye’s role in managing the presence of *T. mercedesae* will be crucial and strategic for beekeeping in the near future. On the other hand, the main problem for a fast reaction is that the mite is classified as a notifiable disease, and notification may result in culling colonies in the affected apiary, followed by quarantine of the apiary area; thus, beekeepers may be reluctant to report suspected cases. As the organism is a parasite, the term “control” rather than “treatment” is more appropriate in accordance with its biology. Rather than relying solely on mandatory notification, emphasis should be placed on epidemiological surveillance and control of its spread, as provisioned in the EU Animal Health Law, according to which measures are needed to prevent Tropilaelaps mites from spreading to parts of the Union, and surveillance should be enacted to ensure early detection in case of arrival (EU, 2018). Thus, the following actions are recommended: systematic monitoring of overwintering areas in the Mediterranean region; establishment of sentinel apiaries; distribution of long-acting, slow-release formic acid preparations to beekeepers through government-supported control programs; increased training efforts on mite detection, transmission pathways and biotechnical control methods such as induced brood interruption, for example via queen caging (Kovačić et al., 2023), and the preparation and distribution of manuals, books, information sheets, protocols for beekeeping best practice. Appropriate official measures should be implemented accordingly. This argues for immediate tripartisan coordination (Georgia–Türkiye–Europe border states), harmonized diagnostic protocols at the points of highest risk (migratory apiaries, conserved genetic reserves, and export-oriented operations), and pre-emptive surveillance particularly in Greece and Bulgaria, given their direct land borders with Türkiye.

## Supporting information

Supp. S1 the morphometric measurements of the samples

## Data availability

The mitochondrial COI sequence was uploaded to GenBank with accession number PZ835872. Voucher specimens are deposited in the Laboratory of MicroBeeotic of Hacettepe University in Ankara.

## Acknowledgements

We would like to thank the Artvin Beekeepers’ Association, especially the President, Mr. İbrahim Durmuş, and Selaaddin Usta, for their collaboration. And also be thankful to Rahime Özçelik and Gülçin İspir, who are beekeeping technicians, for their assistance with all field studies.

## Author contributions

AÖ conducted the field surveillance, detection, and morphometric analysis; IK conducted the molecular analysis; AÖ, AU, CC, and IK equally drafted, edited, reviewed, and approved the final version of the manuscript.

## References

Anderson DL & Roberts JMK (2013) Standard methods for Tropilaelaps mites research, Journal of Apicultural Research, 52:4, 1–16, DOI: 10.3896/IBRA.1.52.4.21

Aurell D, Tokach R, Chuttong B, Praphawilai P, Barascou L, Steury TD, Duffy K, Jung C, Oh H, Bruckner S, Williams GR (2026) First estimates of the population growth rate of the parasitic honey bee mite Tropilaelaps mercedesae in Apis mellifera colonies. bioRxiv 2026.07.06.736813. 10.64898/2026.07.06.736813

Brandorf A, Ivoilova MM, Yañez O, Neumann P, Soroker V (2025) First report of established mite populations, Tropilaelaps mercedesae, in Europe. J Apic Res 64:842–844. 10.1080/00218839.2024.2343976

Chantawannakul, P., Ramsey, S., Vanengelsdorp, D., Khongphinitbunjong, K., & Phokasem, P. (2018). Tropilaelaps mite: An emerging threat to European honey bee. Current Opinion in Insect Science, 26, 69–75. 10.1016/j.cois.2018.01.012

Delfinado, M. D. 1963. Mites of the honey bee in South-East Asia. J Apic Res 2: 113–114

Doyle JJ, Doyle JL (1987) A rapid DNA isolation procedure for small quantities of fresh leaf tissue. Phytochem Bull 19:11–15

Esmaeily M, Kwon S, Babaeian E, Jung C (2026) Tropilaelaps mercedesae: an emerging global threat to apiculture: a comprehensive review. Front Insect Sci 6:1871985. 10.3389/finsc.2026.1871985

EU Commission Implementing Regulation (EU) 2018/1882 of 3 December 2018 on the application of certain disease prevention and control rules to categories of listed diseases and establishing a list of species and groups of species posing a considerable risk for the spread of those listed diseases (http://data.europa.eu/eli/reg_impl/2018/1882/2024-02-01)

Folmer O, Black M, Hoeh W, Lutz R, Vrijenhoek R. DNA primers for amplification of mitochondrial cytochrome c oxidase subunit I from diverse metazoan invertebrates. Mol Mar Biol Biotechnol. 1994 Oct;3(5):294–9.

Franco S, Duquesne V and Laurent M. 2025. “Geographical Spread of the Exotic Mite Tropilaelapsspp.: new reports from the Caucasus”. Scientific Note published on the website of the European Union Reference Laboratory for Bee Health on 15 December 2025. (https://sitesv2.anses.fr/en/system/files/Scientific_Note_EURL_Geograhical_Distribution_Tropilaelaps_December_2025_0.pdf. (Last accessed on 11 September 2026)

Gill M. C., Chuttong B., Davies P., Earl A., Tonge G., & Etheridge D. (2025). In vitro assessment of Tropilaelaps mercedesae survival across different substrates. Apis, 2(2), 22–27. 10.62949/02634299.0581119

Girişgin O, Aydın L, Girişgin AO, Özsoy İA, Topcu D (2026) Economic analysis of Tropilaelaps spp. infestation in honeybees, risk commodities and probabilities for its introduction in Türkiye. Kafkas Univ Vet Fak Derg 32:409–415. 10.9775/kvfd.2026.36466

Goryachev DS, Kuzmich EG (2025) Pests of honey bees in the Belarusian Lakeside Region. In: Proceedings of the international scientific and practical conference for students, MSc students, PhD students and young scientists, Vitebsk State Academy of Veterinary Medicine, Vitebsk, Belarus, 15–16 May 2025

Janashia I, Uzunov A, Chen C, Costa C, Cilia G (2024) First report on Tropilaelaps mercedesae presence in Georgia: the mite is heading westward! J Apic Sci 68:183–188. 10.2478/jas-2024-0010

Joharchi O, Stolbova VV (2024) The first report on the ectoparasitic genus Tropilaelaps Delfinado & Baker (Acari: Mesostigmata: Laelapidae) in Russia. Persian J Acarol 13:513–516. https://www.biotaxa.org/pja/article/view/85545

Kandemir İ, Kence M., Kence A. (2000) Genetic and morphometric variation in honeybee (Apis mellifera L.) populations of Turkey. Apidologie, 31 3 (2000) 343–356. DOI: 10.1051/apido:2000126

Kovačić M, Uzunov A, Tlak Gajger I, Pietropaoli M, Soroker V, Adjlane N, et al (2023) Honey vs. mite: a trade-off strategy by applying summer brood interruption for Varroa destructor control in the Mediterranean region. Insects 14:751. 10.3390/insects14090751

Özkırım A (2018) Beekeeping in Turkey: bridging Asia and Europe. In: Chantawannakul P, Williams G, Neumann P (eds) Asian beekeeping in the 21st century. Springer, Singapore, pp 41–69

Ramirez JL, Tembrock LR, Zink FA, Fife A, Gilligan TM, Chen Y, Evans JD, Mottern J, Smith-Pardo AH and Ochoa R (2026) Interception of an Apis dorsata swarm with Tropilaelaps mercedesae and Kuzinia morsei mites on a cargo vessel inbound to the United States. Front. Insect Sci. 6:1829350. doi: 10.3389/finsc.2026.1829350

Tokach R, Aurell D, Chuttong B, Williams GR (2025) Observation of Tropilaelaps mercedesae (Mesostigmata: Laelapidae) on Western honey bees (Apis mellifera) exiting colonies. J Econ Entomol 118:966–969. 10.1093/jee/toae305

Selis, M., Cilia, G., Wood, T. J., & Soon, V. (2024). Taxonomic revision of the Stenodynerus fastidiosissimus species-group in Western Europe and North Africa (Hymenoptera: Vespidae: Eumeninae), February 2024. Zootaxa, 5418(1), 34–56. 10.11646/zootaxa.5418.1.2

Tokach R, Aurell D, Chuttong B, Williams GR. (2025). Observation of Tropilaelaps mercedesae (Mesostigmata: Laelapidae) on Western honey bees (Apis mellifera) exiting colonies. J Econ Entomol. 2025 Apr 26;118(2):966–969. doi: 10.1093/jee/toae305. PMID: 39888976; PMCID: PMC12034309.

Turkish Statistical Institute. (2025). Livestock production statistics, 2025 [Data set]. www.tuik.gov.tr

Uzunov A, Janashia I, Chen C, Costa C, Kovačić M (2025) A scientific note on ‘rapid brood decapping’: a method for assessment of honey bee (Apis mellifera) brood infestation with Tropilaelaps mercedesae. Apidologie 56:40. 10.1007/s13592-025-01171-2

Uzunov A, Janashia I, Chen C, Costa C, Kovačić M, Gill MC (2026) Swarming promotes Tropilaelaps mercedesae (Mesostigmata: Laelapidae) dispersal in Apis mellifera (Hymenoptera: Apidae). J Econ Entomol 119:1473–1477. 10.1093/jee/toag027

World Organisation for Animal Health (2018) Infestation of honey bees with Tropilaelaps spp. In: Manual of diagnostic tests and vaccines for terrestrial animals, Chapter 3.2.5. Version adopted May 2018. https://www.woah.org/fileadmin/Home/eng/Health_standards/tahm/3.02.05_TROPILAELAPS.pdf

World Organisation for Animal Health (WOAH) (2026) WAHIS event report 7792. World Animal Health Information System. https://wahis.woah.org/#/in-event/7792/dashboard. Accessed 17 August 2026

