## Supplementary figures and images for "First report on *Tropilaelaps mercedesae* presence in Türkiye and insight into potential westward invasion corridors"

### Supp. S1 the morphometric measurements of the samples

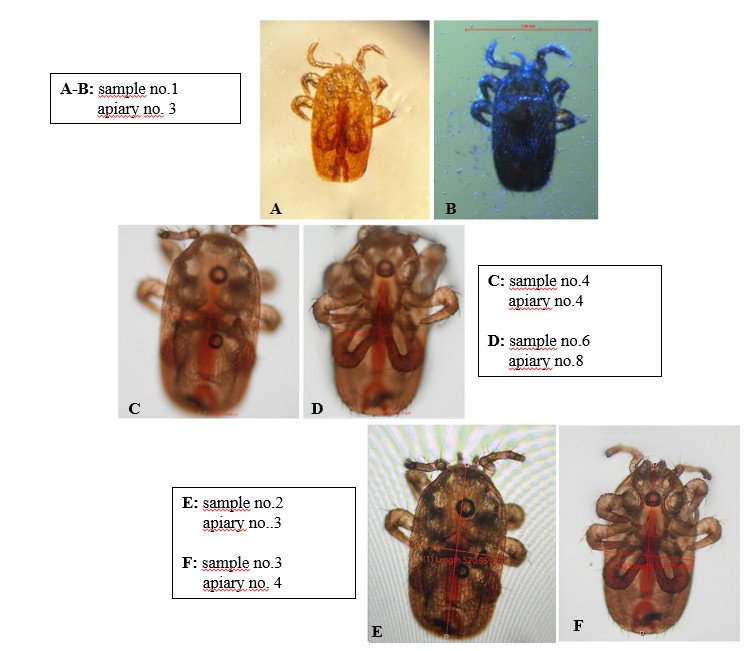
